# Deficient processing of regularity violations during visuospatial neglect: a visual mismatch negativity study

**DOI:** 10.64898/2025.12.10.693186

**Authors:** Nóra Csikós, Petia Kojouharova, Gábor Szabó, Gábor Fazekas, Zsófia Anna Gaál, István Czigler

## Abstract

Visuospatial neglect (VSN) after right-hemisphere stroke causes reduced engagement with left-sided stimuli, but how VSN affects automatic processes, i.e. the registration of regularities and regularity violations is still equivocal. This study investigated these processes using event-related potential components C1, C2/N1, and the visual mismatch negativity (vMMN), and how they differ in VSN compared to stroke and healthy controls, and their changes with time and rehabilitation.

We applied a passive oddball paradigm, where diamond patterns periodically disappeared (OFF events) and reappeared (ON events) on the lower left and right sides, creating two simultaneous but independent sequences with standard and deviant events.

The study included a VSN group (*N*=17, *M*=53.88±0.28 yrs); stroke patients (*N*=16, *M* = 56.81 ± 13.26 yrs); and healthy participants (*N*=17, *M*=53.89±10.79 yrs). The VSN group underwent measurements three times: pre- and post-rehabilitation, and at a 4-month follow-up.

Our results show that the C1 component emerged reliably in healthy participants and as a tendency in stroke controls, but not in VSN at any measurement point. The C2/N1 component emerged for all events without group differences. VSN patients showed no reliable vMMN, while it emerged in stroke patients for left-sided OFF events. Healthy controls showed vMMN in every condition.

Our results suggest that both stroke and VSN impair the automatic detection of regularity violations, but VSN does so in a more substantial way, and this deficit does not improve with time or rehabilitation. Furthermore, the missing C1 component in VSN may indicate an early-stage deficiency even in visual processing.

## Introduction

Unilateral neglect, or in this particular study, visuospatial neglect (VSN), often develops after right-hemisphere damage (Bartolomeo et al., 2007; Husain, 2008; Kerkhoff, 2001). It affects approximately 20-40% of patients with a right hemisphere stroke (Buxbaum et al., 2004; Ringman et al., 2004), and leads to poorer functional recovery compared to stroke alone (Di Monaco et al., 2011; Nijboer et al., 2013). It is often defined as an attentional deficit affecting the contralateral (left) side of space relative to the side of brain damage. Patients fail to orient to, detect, react to, or engage with information coming from the left side of space, even though primary sensory processing and its neural correlates remain intact (Kerkhoff, 2001; Vallar, 1998; Marshall & Robertson, 2013).

Neglect also differs from other syndromes such as hemianopia, which involves a partial loss of the visual field and cannot be ameliorated by cueing to detect stimuli on the affected side. While unilateral neglect can affect any modality, including auditory or sensorimotor domains, VSN refers more specifically to symptoms in the visual modality involving spatial stimuli. Within right-hemisphere stroke, VSN often follows extensive damage in temporal, parietal, or fronto-parietal white matter regions (Bartolomeo et al., 2007, 2012; Corbetta & Shulman, 2011; Ptak & Schnider, 2011), although lesions in other areas such as the basal ganglia or thalamus can also cause VSN (de Schotten et al., 2014; Karnath et al., 2002).

Most studies on neglect have focused on attention-related deficits as the underlying mechanism of this syndrome. Although findings still remain equivocal, there is consensus that the symptoms arise from widespread functional disruption both within and between the dorsal, goal-directed (DAN), and ventral, stimulus-driven (VAN) large-scale attentional networks, as well as their interconnecting white matter pathways (Corbetta & Shulman, 2011; Bartolomeo et al., 2012). Among these, the VAN is strongly lateralised to the right hemisphere and encompasses regions such as the temporoparietal junction and middle frontal gyri, with the superior longitudinal fasciculus III (SLF III) serving as the principal white matter tract connecting parietal and frontal components of the VAN (Bartolomeo et al., 2012). This network is often the one directly compromised by the structural (and thus, functional) damage caused by stroke, resulting in deficits in bottom-up, exogenous orienting of attention, even towards salient and behaviourally relevant events within the left visual field (Karnath, 2015). Additionally, studies have found that SLF II white matter tracts, which provide intrahemispheric connections between the VAN and DAN, can be frequently structurally damaged, leading to disconnection between these two attentional systems (De Schotten et al., 2014). This disconnection may also contribute to impaired functioning within the DAN, even where it remains structurally intact within the affected right hemisphere (Bartolomeo et al., 2007; 2012). The DAN is bilaterally represented and comprises frontal and parietal regions, including the intraparietal sulcus and frontal eye fields, connected via the SLF I. Reduced functional activity within this network (on the right) can disrupt top-down control of endogenous spatial orienting and feedback modulation of early sensory processes. This right-hemispheric hypoactivation may be accompanied by hyperactivation of the left DAN, contributing to the rightward attentional bias characteristic of the syndrome (Vuilleumier, 2013).

These deficits in spatial attentional functions can be augmented by impairments in non-spatial processes such as arousal and alertness. Reduced phasic and tonic arousal negatively influence attentional functions, as patients struggle to adequately recruit these systems in response to spatial stimuli, particularly those appearing on the left (Robertson et al., 1998; DeGutis & Van Vleet, 2010; Chica et al., 2012). Such impairments may further limit compensatory mechanisms, resulting in more adverse outcomes than those associated with purely spatial deficits (Corbetta & Shulman, 2011; Bartolomeo et al, 2012). Nevertheless, altered arousal and alertness are not exclusive symptoms of neglect, they are often characteristic to right-hemisphere stroke without neglect as well (Spaccavento et al., 2019; Villarreal et al., 2021).

Overall, the literature predominantly conceptualises neglect as an attentional disorder. However, research so far largely overlooked the potential role of earlier, nonattentive stages of sensory processing. To address this gap, the present study aims to investigate non-attentive visual processing by examining ERP components such as C1, C2 and the visual mismatch negativity (vMMN). This approach enables the exploration of whether early, automatic sensory discrimination processes are affected and form a more comprehensive understanding of neural impairments associated with neglect.

Primary sensory processing is generally regarded as intact in neglect. Electrophysiological evidence supporting this view derives from event-related potential (ERP) studies investigating exogenous components such as C1, P1, and C2. Traditionally, these components are interpreted as signatures of subsequent steps of processing visual stimuli: C1, emerging at approximately 60–80 ms post-stimulus, reflects the earliest stages of visual processing and is particularly sensitive to features such as contrast and contour; C2, appearing around 100-120 ms, is associated with the further integration and more detail-oriented processing of incoming stimuli; and C3, typically observed around 140-160 ms, is indicative of more complex cognitive processing involving attention and memory (Luck, 2012). For instance, DiRusso et al. (2008), found that in unilateral neglect patients, ERP components such as C1 for left-sided stimuli remained unaltered up to appr. 130 ms post-stimulus, showing patterns comparable with right-sided stimuli and healthy controls. While C2 and C3 have not yet been directly examined in visual neglect, based on Di Russo’s findings, where deficits emerged only after 130 ms, these components being more closely related to attention-dependent mechanisms, are more likely to be affected.

Other automatic, but more complex processes involved in visuospatial event processing include (1) the ability to extract regularities from a series of stimuli, and develop rules to predict the forthcoming sequence of stimuli; and (2) the ability to detect and register rare or unexpected events that violate these regularities. The latter process, automatic change detection, is reflected in an ERP component, the visual mismatch negativity (vMMN) (Stefanics et al., 2011), the counterpart of the more frequently studied auditory MMN, that usually emerges between 100-300 ms post-stimulus with negative polarity (for reviews see Stefanics et al., 201; Czigler & Kojouharova, 2021). VMMN is commonly investigated using a passive oddball paradigm, where sequences of frequently presented stimuli (standard events) are occasionally interrupted by different, infrequently presented stimuli (deviant events). Since neither the standards nor the deviants are relevant or related to the ongoing task, this paradigm is considered ‘passive’. VMMN can also be elicited using a wide variety of visual stimuli, e.g. such as color (Athanasopoulos et al., 2010), spatial frequency (Heslenfeld, 2005), movement direction (Lorenzo-López et al., 2004), and orientation (Kimura et al., 2009). VMMN is also elicited by deviant higher-order visual characteristics, like object-related deviancy (Müller et al., 2010), facial attributes (Astikainen & Hietanen, 2009; Kreegipuu et al., 2013; Kecskés-Kovács et al., 2013), and semantic categories (Hu et al., 2020). This wide variety of results demonstrates the vMMN is a reliable indicator of violation of established sequential regularities in the visual modality. Moreover, it has been applied in a variety of clinically oriented investigations across diverse populations.

VMMN is frequently interpreted within the framework of ‘predictive coding theory’ (Garrido et al., 2008, Stefanics et al., 2014). According to this theory, vMMN reflects a cascade of comparison processes between the memory-based expectations of events and the representations of the actual incoming sensory events. This cascade begins when an unexpected event fails to conform to internal expectations, and thus producing a mismatch referred to as a ‘prediction error’. This cascade then concludes when the updated memory content matches with the incoming sensory representation. This view suggests that vMMN depends on the properties of memory content established by regular event sequences, and is elicited when a change in that sequence is registered as a prediction error, thereby triggering this cascade-like processing.

VMMN may serve as a valuable tool for understanding how automatic change detection, an integral component of the ability to form and predict regularities, is altered in VSN patients. To our knowledge, only two studies to date have investigated automatic change detection in neglect, and both studies applied auditory mismatch paradigms to examine aMMN. The first by Deouell et al. (2000) found that the automatic detection of spatial deviancies in sounds were impaired for left-side stimulation, but remained intact for right-side stimulation. Another important study by Doricchi et al. (2021) employed MMN (and P3) within an auditory oddball paradigm using sequences of tones presented central to the participants, where deviants could occur on the left, the right, or as omissions relative to the standards. Their results showed that left-side deviants failed to elicit aMMN, whereas right-side deviants did, thus supporting Deouell’s earlier findings. Altogether, these two studies suggest that, at least in the auditory domain, the automatic registration of regularity violations is deficient on the left side. This deficiency could reflect a disruption in the cascade-like process of testing and updating internal predictions, or a failure to detect stimuli that might violate or update these regularities. Nonetheless, so far there has been no study investigating vMMN in VSN, even though visual deficits in neglect are more common and frequently more severe than auditory ones (Gainotti, 2010; Stone et al., 1993).

The aim of the present study was to examine the C1, C2/N1 and vMMN components in order to determine how automatic processes of event detection and the registration of regularity violations in the visual modality are affected by VSN, in comparison with right-hemisphere stroke without neglect and healthy individuals together with how those may change over time and with rehabilitation. To be able to investigate these ERP components separately for left- and right-side stimuli presentation, we applied a passive oddball paradigm where we presented two independent, but simultaneously appearing stimuli sequences to the lower left and right side of the screen (Csikós et al., 2023). This was important, as in everyday life, VSN patients, just as healthy individuals are typically exposed to multiple concurrent streams of stimuli sequences, or to information coming from both of their sides at the same time. This task was specifically developed to enable us to examine the event registration and change detection processes separately for the two sides, reflecting the fact that VSN is often characterised by relatively preserved perception on the right, but deficits on the left.

We adopted the OFF-ON method of stimulation (Sulykos et al., 2017), where the events, geometric diamond patterns in this case, are formed by certain parts of the diamonds disappearing (OFF events) and then reappearing again (ON events), creating two distinct type of events that could be processed by the cognitive system. This method of stimulation is particularly useful, prolonged presentation of the complete shapes during ON events promotes low-level adaptation; thus, the contrast between standard and deviant events is less confounded by adaptation of low-level input structures. Previous studies using this method found that in case of non-lateralized stimuli arrangement and a single sequence of events, vMMN emerged only to OFF events, presumably because the reappearance of the whole shapes (ON events) did not count as ‘surprising’ (Sulykos et al., 2017). Another study with the same stimuli arrangement as here (Csikós et al., 2023) found that participants were able to adequately process the stimuli sequences from both sides, signaled by the emergence of exogenous components, but not to their full extent, and as a result only produced vMNN for OFF events from left-side stimulation, and for ON events from right-side stimulation.

We recruited three groups: patients with right-hemisphere stroke who developed VSN, stroke patients without VSN, and healthy controls. VSN and stroke patients during the testing were undergoing inpatient rehabilitation, and the VSN group was assessed at three time points: at the start and at the end of their rehabilitation, and appr. four months after. This longitudinal design allowed us not only to examine how automatic change detection functions in VSN patients, but also whether it evolves over time with recovery. VSN is traditionally described as a syndrome affecting the left side of space, but the results are equivocal about how intact implicit processing is on that side. Thus, our main question was if vMMN to left-side stimulation would be attenuated, missing, or remain intact, whereas we hypothesized that it would be present to right-side stimulation, comparable to that of healthy controls. Moreover, we also expected the exogenous components C1 and C2/N1 to appear to both left-and right-side stimulation without deficits. Additionally, we aimed to investigate if VSN can have a more pronounced negative effect than stroke only. Lastly, we explored if C1, C2/N1 and vMMN in the VSN group to either side of stimulation change over time, and if so, in what manner.

## Methods

### Participants

There were 50 participants altogether, of which 17 were VSN patients (11 female, *M* = 53.88 ± 10.28 yrs), 16 were age-matched stroke patients without VSN (8 female, *M* = 56.81 ± 13.26 yrs), and 17 were healthy age-matched control participants (7 female, *M* = 53.89 ± 10.79 yrs). Due to abundant alpha activity, an extreme number of artefacts or their age not being a match, the results of 5 participants out of the original 21 were discarded in the Stroke control group, and 7 out of the original 24 in the Healthy control group. Participants had normal or corrected to normal vision (measured with Snellen cards). The study was conducted in accordance with the Declaration of Helsinki, and approved by the Medical Research Council, Research Ethics Committee of Hungary (TUKEB). Written informed consent was obtained from all participants.

VSN and stroke patients were examined and rehabilitated at the Rehabilitation Clinic of Semmelweis University, Hungary. Patients with bilateral strokes, signs of dementia measured by the Mini Mental State Examination (MMSE) test, hemianopia, or any preexisting neurological disorders were excluded from the study. At the time of the data collection, all patients were free from confusion and temporal or spatial disorientation. Both these groups consisted of patients with only right hemisphere strokes described in Table 1. Patients were extensively screened by neuropsychologists at the hospital for VSN before recruiting them into this study, and all participants in the VSN group were given the diagnosis of hemineglect. In addition to that, VSN was assessed with two neuropsychological tests within the experiment in the two patient groups, the Line Bisection test (McIntosh et al., 2017; Schenkenberg et al., 1980) and the Bells test (Gauthier et al., 1989). T-tests were calculated to show the differences on these two tests between the patient groups: for both tests (t(31) = 3.40, p = 0.001 for LB and t(31) = 2.70, p = 0.01 for the Bells test) the difference was significant, showing worse performance in the VSN group. Table 1 contains the demographic, clinical, and neuropsychological data for the VSN, Stroke and Healthy groups.

**Table 1.**
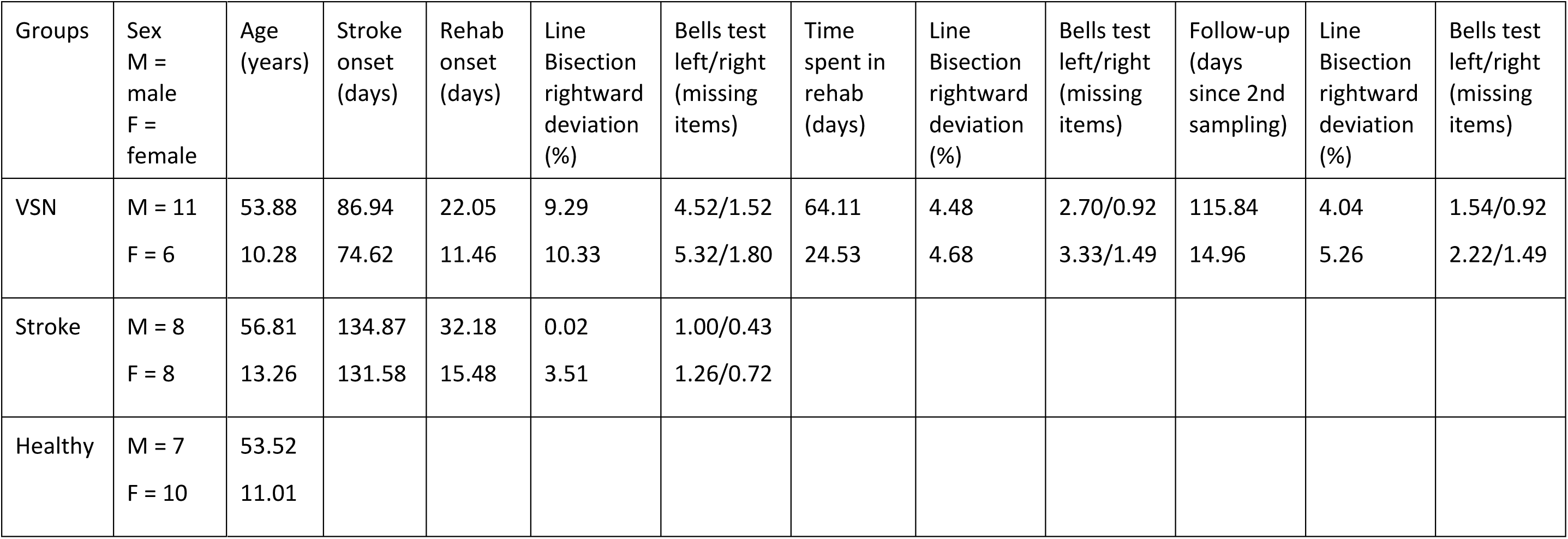
Mean values of demographic, clinical, and neuropsychological data in the visuospatial neglect (VSN), stroke control, and healthy control groups. Stroke onset and Rehabilitation onset were calculated as the number of days relative to the time of the first data sampling. Time spent in rehabilitation indicates the number of consecutive days in in-patient rehabilitation at the time of the second data sampling. (Mean: upper row; SD: lower row)

### Stimuli and procedure

Task and task-relevant stimulus: The stimuli were presented on a 19-inch LCD monitor with a 60 Hz refresh rate. Participants were instructed to perform a reaction-time task, which was embedded into a passive oddball paradigm. The task-relevant stimulus was a centrally located fixation cross that periodically changed its shape, with the unequal vertical and horizontal lines (0.62° and 1.08°) being reversed randomly between 5 and 15 s. Participants were instructed to fixate on this central cross, avoid looking elsewhere during the task, and press the SPACE bar on the keyboard as fast as possible whenever they detected the task-relevant stimulus change. This exercise was made to prevent them from allocating their attention to the vMMN-related peripheral stimuli. Behavioral data were defined as the average reaction-time (RT) and the number of missing detections of the changes.

Task-irrelevant stimuli: The task-irrelevant, vMMN-related stimuli were diamond patterns with their diagonals placed in the lower left and right side of the screen. During the task, either the 45° or 135° parallel lines of the diamonds disappeared for 400 ms +/− 16.6 ms (OFF events) and then reappeared again for 400 ms +/− 16.6 ms (ON events), but the probability of which pair of the parallel lines would vanish were biased, thus creating frequent events (standards) and rare events (deviants). There were no inter-stimulus intervals. These lateral event sequences on the left and right side were presented simultaneously, but they were independent of each other. The standard events of one side were the deviant on the other, and vice versa. In this way we created for different conditions in which vMMN could emerge: OFF events presented on the left side (Left OFF); OFF events presented on the right side (Right OFF); ON events presented on the left side (Left ON); ON events presented on the right side (Right ON). Within a session there were 6 blocks, each consisting of 100 changes from whole diamonds to diamonds with vanished sides, from which 85% were standards, 15% were deviants, in each of the event sequences.

Fig. 1 illustrates the vMMN-related stimuli, and the event sequences. From the 0.9 m viewing distance the size of a diamond was 2.69°. The distance of the diamonds from the imaginary vertical midline of the screen was 3.99°. The vertical distance of the centers of diamonds was 3.67°. The line thickness of the diamonds was 10 pixels. Luminance of the screen and the lines of the diamonds were 45 and 175 cd/m^2^, respectively.

**Fig. 1.**
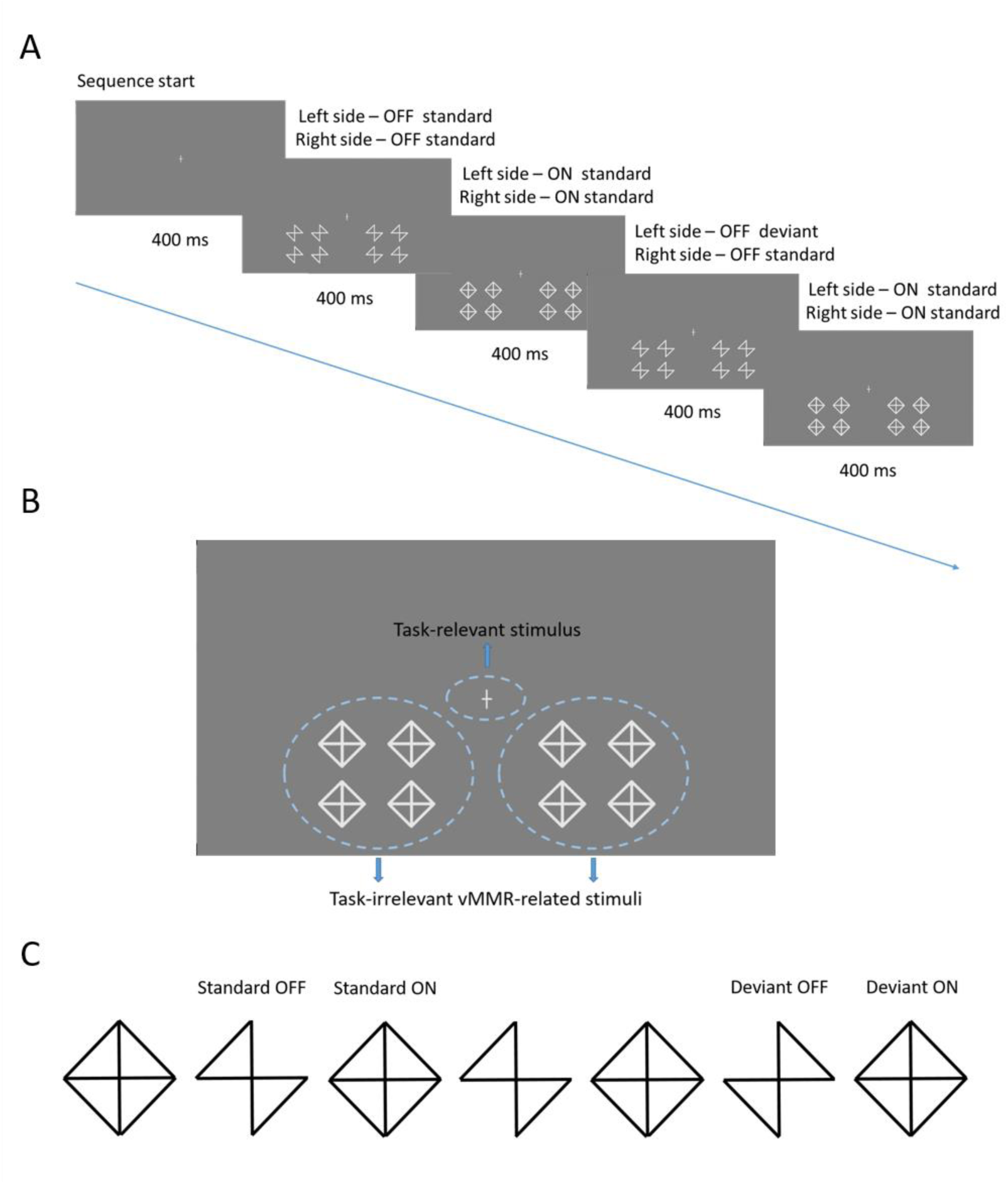
A: Stimuli presented on the lower left and right sides of the visual field at the same time, but within independent sequences with a central fixation cross. The standards on the left side were the deviants on the right side, and vice versa. Both ON and OFF events were presented for 400 ms. B: The display of stimuli on the screen. The central fixation cross is the task-relevant stimulus, the diamond patterns are the task-irrelevant vMMR-related stimuli. C: Examples of how OFF and ON events follow each other in the sequences of vMMR-related stimuli.

In this experiment, the VSN, Stroke, and Healthy control groups had slightly different sampling protocols. The task described above for measuring ERPs were the same across all three groups. Additionally, patients in the VSN group underwent testing at three different time points: at the beginning of their inpatient rehabilitation program, at the end of the program, and approximately at a four-month follow-up. Each time, after their EEG was measured, they were assessed using the Line Bisection and Bells Test neuropsychological tests. Patients in the Stroke group were assessed only once but completed both the ERP-related task and the neuropsychological tests. Healthy control participants only completed the ERP-related task while measuring their EEG.

### Measurement of electrical brain activity

Electrical brain activity was recorded from 10 locations according to the extended 10-20 system (F3, F4, PO7, PO3, POZ, PO4, PO8, O1, OZ, O2) with NuAmps amplifiers (bandpass: DC-70 Hz) using NeuroScan 4.4 software (sampling rate: 1000 Hz) and Ag/AgCl active electrodes. The reference electrode was placed on the nose tip, and the ground electrode on the forehead (AFz). Horizontal and vertical electrooculograms (HEOG and VEOG) were recorded with bipolar configurations between two electrodes (placed lateral to the outer canthi of the two eyes and above and below the left eye, respectively). The EEG signal was bandpass filtered offline with a non-causal Kaiser-windowed Finite Impulse Response filter (low pass filter parameters: 30 Hz of cutoff frequency, beta of 12.265, a transition bandwidth of 10 Hz; high pass filter parameters: 0.1 Hz of cutoff frequency). The EEG data were processed with MATLAB R2015a (version 2015a, MathWorks, Natick, MA). Independent component analysis (ICA) was used to remove eye-movement artefacts. Epochs with larger than 100 μV voltage change between the minimum and maximum of the epoch at any electrode site were considered artifacts and rejected from further processing. Table 2 shows the mean number of averaged epochs for each event.

**Table 2.** Mean number of standard and deviant epochs averaged together for the Left OFF, Left ON, Right OFF, Right ON events (S.M.E. in parenthesis).

| Condition | Left OFF |  | Left ON |  | Right OFF |  | Right ON |  |
| --- | --- | --- | --- | --- | --- | --- | --- | --- |
| Event | Standard | Deviant | Standard | Deviant | Standard | Deviant | Standard | Deviant |
| VSN 1 | 75.82<br>(7.82) | 76.20<br>(7.35) | 76.02<br>(6.99) | 76.26<br>(6.49) | 76.00<br>(5.89) | 76.08<br>(6.95) | 75.29<br>(7.23) | 75.50<br>(7.32) |
| VSN 2 | 77.20<br>(6.39) | 77.82<br>(4.93) | 76.91<br>(6.25) | 77.80<br>(5.11) | 77.73<br>(5.00) | 77.55<br>(6.44) | 77.91<br>(5.14) | 78.02<br>(6.24) |
| VSN 3 | 79.65<br>(3.08) | 80.30<br>(3.69) | 79.38<br>(3.16) | 80.19<br>(4.63) | 80.46<br>(3.69) | 79.23<br>(3.92) | 80.19<br>(4.64) | 79.00<br>(3.87) |
| Stroke | 78.93<br>(4.71) | 77.38<br>(5.79) | 78.75<br>(4.94) | 76.96<br>(5.74) | 78.84<br>(4.40) | 78.31<br>(5.58) | 79.15<br>(4.29) | 77.96<br>(5.67) |
| Healthy | 79.81<br>(2.78) | 79.75<br>(3.57) | 79.53<br>(2.75) | 79.34<br>(3.04) | 79.25<br>(3.76) | 79.56<br>(3.63) | 79.28<br>(3.25) | 79.93<br>(2.81) |

Stimulus onset was measured by a photodiode, providing an exact zero value for averaging. Epochs were extracted for further analysis ranging from −100 to 400 ms for both the ON and OFF events from both the left and right side, then we averaged them separately. In order to have roughly the same number of epochs for standard and deviant events, we only calculated with standard events preceding a deviant. The first 100 ms ranging from −100 to 0 relative to stimulus presentation served as the baseline.

To measure the amplitude and latency of the exogenous components (C1 and C2) and the emergence of vMMN, we constructed left- and a right-side ROIs from the PO3, O1, and PO4, O2 locations based on our previous studies (File et al., 2017; Csikós et al., 2023).

Peak amplitude values of the exogenous components were measured as the +/− 10 ms range around the amplitude maxima. Then, one-sample t-tests were calculated in every group, and where relevant, the results were compared in Repeated Measures of ANOVAs with factors of *Group* (VSN 1, Stroke, Healthy)*, Side* (left-, right-side stimulation), *Event* (OFF, ON), *ROI* (right, left).

To investigate vMMN, we calculated difference potentials from the amplitude values in the two ROIs. While the same stimulus configuration appeared as deviant in a sequence and as standard in another sequence, difference potentials were calculated from physically identical ERPs. Considering the results of previous studies (Sulykos et al., 2017; Czigler et al., 2019; Csikós et al., 2023), we expected vMMN to emerge in a range of approximately 90-180 ms. As a first step, we conducted one-sample t-tests within this range separately for every condition. We considered vMMN to emerge if the difference was significant (at least 20 consecutive values, equivalent to 20 ms) within a ROI at a significance level of p<0.05. Based on these calculations, we obtained a time range of 110-140 ms for the Right OFF condition, and 90-120 ms for the Left OFF, Left ON, Right ON conditions.

As a second step, we calculated t-tests in every group for every condition in those time ranges, and where relevant, we followed up with ANOVAs with factors of *Side* (left, right), *Stimulus* (OFF, ON) and *ROI* (right, left). For post hoc paired comparison we used Tukey HSD tests. In these tests the significance level was at least p<0.05. We only present the results where the significant main effect of *Stimulus* and interactions involving the *Stimulus* factor were present. Other effects are available on request. We used the Statistica package (Version 13.4.0.14, TIBCO Software Inc.) for statistical analyses.

## Results

### Behavioral results

#### VSN group

##### Pre-rehabilitation

The average hit rate for the change of fixation was 88.80% (SD = 12.53); the average RT was 660.42 ms (SD = 288.65), respectively. In t-tests, there were no significant differences between the left vs. right side oddball deviancy conditions (hit rate: t(17) = −0.26, p = 0.789; RT: t(17) = 1.00, p = 0.323).

##### Post-rehabilitation

The average hit rate for the change of fixation was 93.24% (SD = 4.23); the average RT was 622.51 ms (SD = 210.56), respectively. In t-tests, there were no significant differences between the left vs. right side oddball deviancy conditions (hit rate: t(17) = 0.23, p = 0.813; RT: t(17) = −0.18, p = 0.851).

##### Follow-up

The average hit rate for the change of fixation was 94.35% (SD = 4.38); the average RT was 574.91 ms (SD = 180.73), respectively. In t-tests, there were no significant differences between the left vs. right side oddball deviancy conditions (hit rate: t(13) = 0.99, p = 0.329; RT: t(13) = 0.19, p = 0.849).

##### Within group comparison

The difference in hit rate between the three data sampling times in the VSN group was significant (2,24)=3.64, p=0.041, η^p2^=0.23), measured by ANOVAs. According to the Tukey HSD post-hoc test, their performance was significantly better at follow-up than at pre-rehabilitation. There was no significant difference in RT.

#### Stroke control group

The average hit rate for the change of fixation was 95.75% (SD = 2.51); the average RT was 514.73 ms (SD = 86.82), respectively. In t-tests, there were no significant differences between the left vs. right side oddball deviancy conditions (hit rate: t(16) = −0.06, p = 0.944; RT: t(16) = −0.40, p = 0.685).

#### Healthy control group

The average hit rate for the change of fixation was 96.59% (SD = 2.22); the average RT was 443.86 ms (SD = 63.46), respectively. In t-tests, there were no significant differences between the left vs. right side oddball deviancy conditions (hit rate: t(17) = 1.19, p = 0.239; RT: t(17) = −0.37, p = 0.709).

#### Group comparisons

The difference in hit rate between the VSN group at pre-rehabilitation and the two control groups was significant (F(2,30)=6.72, p=0.003, η_p_^2^=0.30), measured by ANOVAs. According to the Tukey HSD post-hoc test, the VSN group performed significantly worse than both the Stroke (p = 0.010) and Healthy control groups (p = 0.006), while there was no significant difference between the two control groups. The difference in hit rate remained significant when comparing the VSN group at post-rehabilitation with the two control groups (F(2,30)=4.59, p=0.018, η_p_^2^=0.23), but the difference was no longer significant when comparing the VSN group at follow-up, respectively.

The difference in RT between the VSN group at pre-rehabilitation and the two control groups was significant (F(2,30)=6.73, p=0.003, η_p_^2^=0.30), measured by ANOVAs. According to the Tukey HSD post-hoc test, the VSN group was significantly slower than the Healthy control group (p = 0.003), and the difference between the VSN and Stroke group approached significance (p = 0.060). There was no significant difference between the two control groups. That did not change, the difference in RT was significant when comparing the VSN group at post-rehabilitation (F(2,30)=7.63, p=0.002, η_p_^2^=0.25), and then at follow-up (F(2,24)=4.05, p=0.030, η_p_^2^=0.25) with the two control groups, respectively.

### Event-related potentials

#### Exogenous components

Table 3 shows the mean amplitude values of the exogenous components C1 and C2 in the Left OFF, Left ON, Right OFF, and Right ON conditions at the left and right ROIs for all three groups. As Figure 2 shows, C1 was elicited only by the ON events. According to the t-tests, in the Healthy control group, this component emerged at both ROIs for left-side stimulation (t(16)=3.88, p=0.001; t(16)= 3.85, p=0.001) and at the left ROI for right-side stimulation (t(16)= 2.55, p<0.022; Benjamini-Hochberg adjusted values). In the Stroke control group, we obtained near-significant values at the left ROI for both left- and right-side stimulation (t(15)=2.45, p=0.066; t(15)=2.25, p=0.066; Benjamini-Hochberg adjusted values). In the VSN group reliable C1 emerged only in the 2nd session (post-rehabilitation) for right-side simulation at the left ROI (t16)=2.35, p=0.031), whereas for right-side stimulation at the right ROI (t(16)=2.07, p=0.055) approached significance (Benjamini-Hochberg adjusted values).

**Table 3.** T-test values for the C2 exogenous component in the Left OFF, Left ON, Right OFF, Right ON conditions at the left and right ROIs, separately for all groups (Bejamini-Hochberg adjusted values).

| Condition | Left OFF |  | Left ON |  | Right OFF |  | Right ON |  |
| --- | --- | --- | --- | --- | --- | --- | --- | --- |
|  | Left | Right | Left | Right | Left | Right | Left | Right |
| VSN 1<br>(N=17) | -3.63 | -2.65 | -5.03 | -3.62 | -3.39 | -2.21 | -5.46 | -5.29 |
| VSN 2<br>(N=17) | -2.99 | -2.91 | -3.71 | -3.76 | -4.79 | -4.20 | -4.64 | -4.55 |
| VSN 3<br>(N=13) | -3.11 | -2.47 | -5.66 | -5.29 | -2.47 | -1.65<sub>(n.s.)</sub> | -3.85 | -3.59 |
| Stroke<br>(N=16) | -3.68 | -4.00 | -3.96 | -3.94 | -3.67 | -4.16 | -4.88 | -4.53 |
| Healthy<br>(N=17) | -6.10 | -4.60 | -6.27 | -4.95 | -6.99 | -5.67 | -10.66 | -7.30 |

n.s.: The t-test was not significant.

**Fig. 2.**
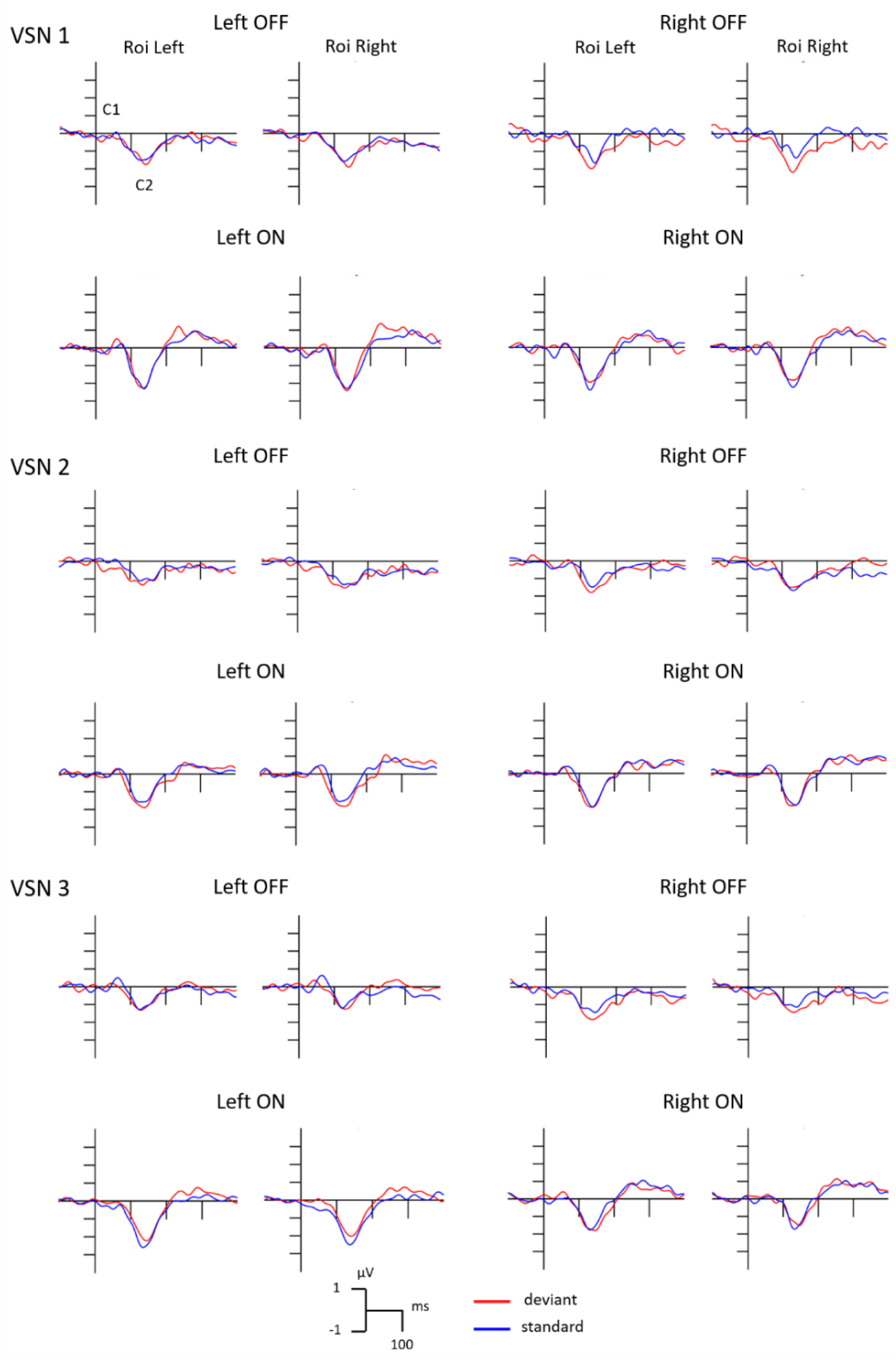
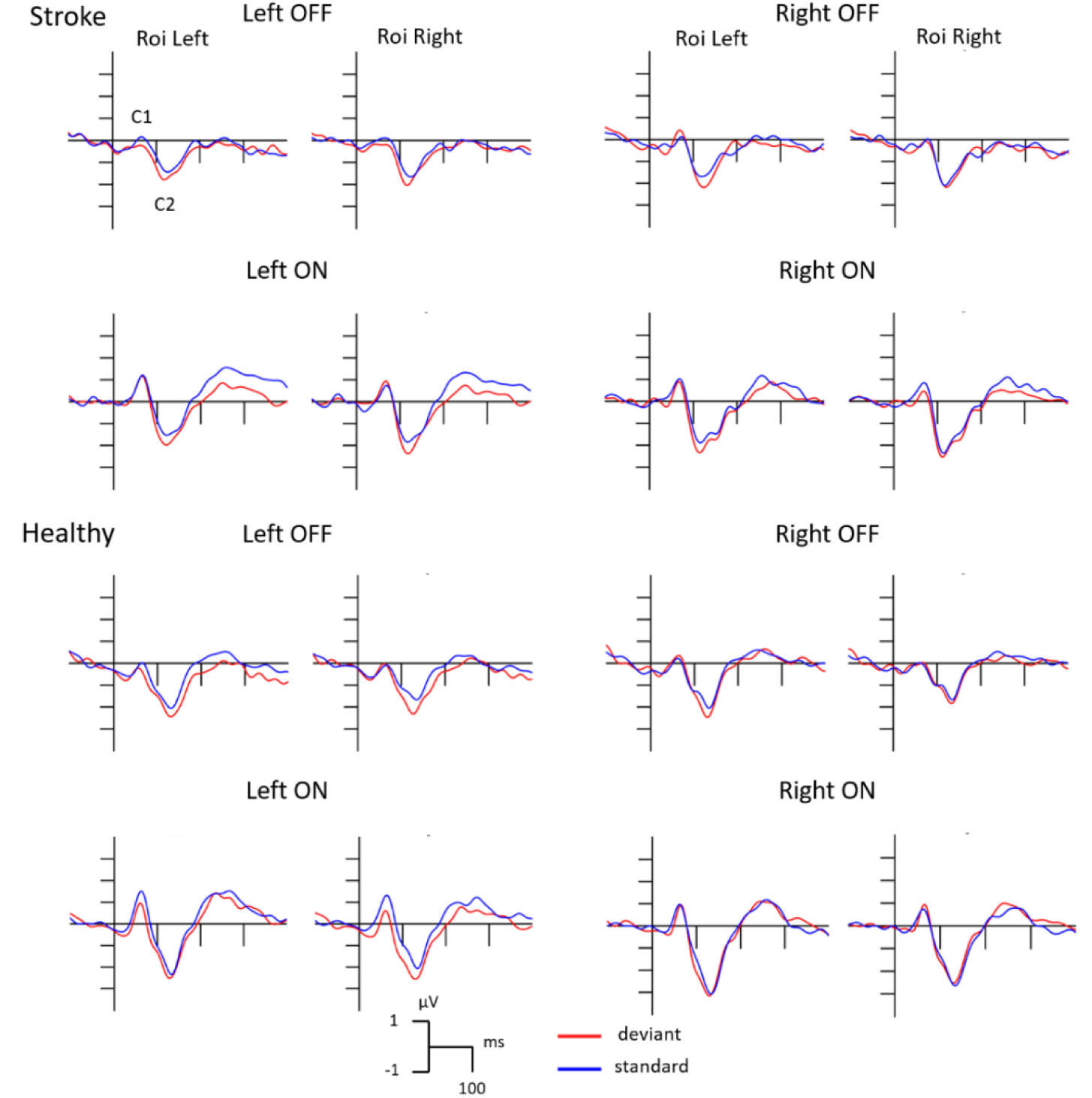
Event-related potentials to standard and deviant stimuli separately for the Left OFF, Left ON, Right OFF, Right ON conditions at the left and right ROIs in all groups and at all measurements in the VSN group.

Figure 2 shows that the second exogenous component (C2/N1) appeared in all groups, in all but one case (VSN group, 3rd session, OFF event, right-side stimulation, right ROI), the amplitude was different from zero as can be seen in Table 4 showing the results of the t-tests. In order to compare the appearance of the C2 component in each group, we conducted Repeated Measures ANOVAs with factors of *Group* (VSN 1, Stroke, Healthy) *x Side* (left-, right-side stimulation) x *Event* (OFF, ON) x *ROI* (right, left). Based on these, the main effect of Event (F(1,47)=8.48, p=0.005, η_p_^2^=0.15), and the *Group x Side x ROI* interaction (F(2,47)=3.35, p=0.004, η_p_^2^=0.12) were significant, respectively, but the post-hoc test showed there were no differences between the groups.

**Table 4.** Mean amplitude values (µV) of the C1 and C2 components to the standards of the Left OFF, Left ON, Right OFF, Right ON conditions at the left and right ROIs, separately for all groups (S.M.E. in parenthesis).

| Comp-<br>onent | Condition | Left OFF |  | Left ON |  | Right OFF |  | Right ON |  |
| --- | --- | --- | --- | --- | --- | --- | --- | --- | --- |
|  | ROI | Left | Right | Left | Right | Left | Right | Left | Right |
| C1 | VSN 1 | -0.04<br>(0.98) | 0.01<br>(1.10) | 0.15<br>(0.86) | -0.07<br>(0.72) | 0.04<br>(1.24) | 0.11<br>(1.34) | 0.17<br>(0.67) | 0.19<br>(0.47) |
|  | VSN 2 | 0.05<br>(0.79) | -0.10<br>(0.87) | 0.25<br>(0.84) | 0.46<br>(0.84) | -0.13<br>(0.64) | -0.45<br>(0.89) | 0.37<br>(0.65) | 0.43<br>(0.87) |
|  | VSN 3 | 0.43<br>(0.38) | 0.57<br>(0.72) | -0.27<br>(0.86) | -0.56<br>(0.97) | -0.19<br>(0.89) | -0.07<br>(1.05) | 0.04<br>(0.62) | 0.12<br>(0.82) |
|  | Stroke | 0.10<br>(0.87) | 0.09<br>(0.99) | 1.07<br>(1.78) | 0.63<br>(1.89) | -0.07<br>(0.58) | -0.17<br>(0.69) | 0.91<br>(1.48) | 0.76<br>(1.74) |
|  | Healthy | -0.09<br>(0.65) | -0.17<br>(0.47) | 1.36<br>(1.45) | 1.19<br>(1.26) | 0.14<br>(0.63) | -0.01<br>(0.69) | 0.81<br>(1.31) | 0.62<br>(1.26) |
| C2 | VSN 1 | -1.49<br>(1.70) | -1.54<br>(2.39) | -2.20<br>(1.80) | -2.22<br>(2.52) | -1.54<br>(1.87) | -1.28<br>(2.40) | -2.29<br>(1.73) | -2.17<br>(1.69) |
|  | VSN 2 | -1.06<br>(1.47) | -1.28<br>(1.81) | -1.56<br>(1.73) | -1.52<br>(1.67) | -1.43<br>(1.23) | -1.64<br>(1.61) | -1.82<br>(1.61) | -1.75<br>(1.58) |
|  | VSN 3 | -1.22<br>(1.41) | -1.12<br>(1.64) | -2.53<br>(1.61) | -2.44<br>(1.66) | -1.40<br>(2.03) | -1.09<br>(2.38) | -1.71<br>(1.60) | -1.64<br>(1.65) |
|  | Stroke | -1.39<br>(1.51) | -1.61<br>(1.61) | -1.47<br>(1.48) | -1.78<br>(1.81) | -1.65<br>(1.80) | -2.01<br>(1.93) | -1.80<br>(1.47) | -2.25<br>(1.99) |
|  | Healthy | -1.97<br>(1.33) | -1.60<br>(1.43) | -2.22<br>(1.46) | -1.95<br>(1.62) | -1.95<br>(1.15) | -1.56<br>(1.13) | -2.97<br>(1.14) | -2.54<br>(1.43) |

Figure 2 shows the ERPs to standard and deviant stimuli in all groups to OFF and ON events at the left and right ROIs. Table 4 shows the mean amplitude values of the exogenous components C1 and C2 in response to the OFF and ON events at both ROIs in all groups.

#### Visual mismatch negativity

The deviant minus standard difference potentials are shown in Fig. 3. Table 5 shows the mean amplitude values of the difference potentials in the Left OFF, Left ON, Right OFF, and Right ON conditions at the left and right ROIs for all three groups. As a note, in the case of VSN at follow-up (VSN 3), the number of participants did not reach the quota needed to reach reliable effect size (based on our G*power analysis), although the analyses were run, the results should be treated with more caution.

**Fig. 3.**
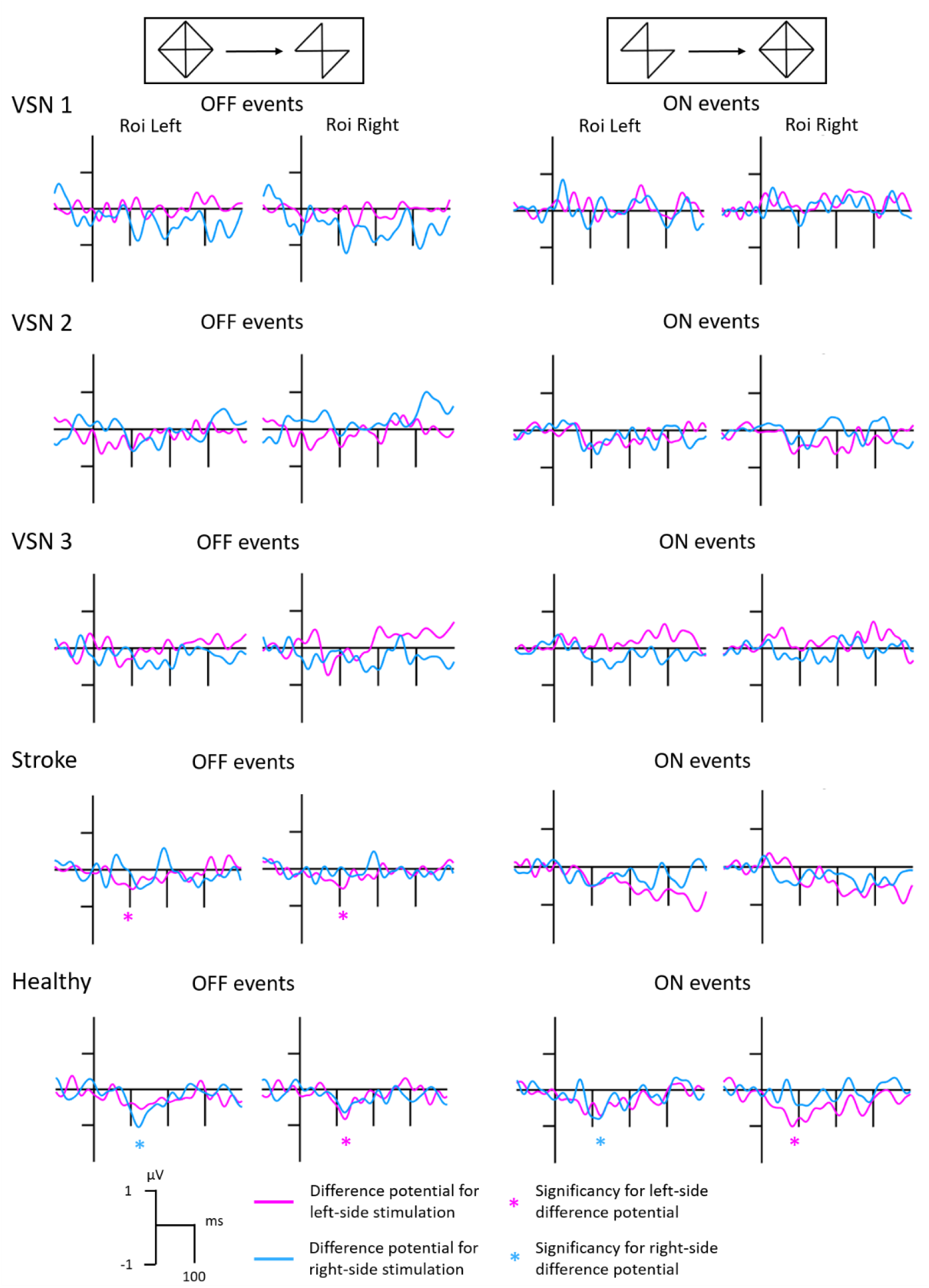
Difference potentials separately for the OFF and ON events from left- and right-side stimulation at the left and right ROIs in all groups, with significant t-test results marked.

**Table 5.** Mean amplitude values (µV) of the difference potentials in the Left OFF, Left ON, Right OFF, Right ON conditions at the left and right ROIs, separately for all groups (S.M.E. in parenthesis).

| Condition | Left OFF |  | Left ON |  | Right OFF |  | Right ON |  |
| --- | --- | --- | --- | --- | --- | --- | --- | --- |
| ROI | Left | Right | Left | Right | Left | Right | Left | Right |
| VSN 1 | 0.11<br>(1.15) | -0.03<br>(0.99) | 0.07<br>(1.46) | 0.04<br>(1.70) | -0.70<br>(1.48) | -1.01<br>(1.56) | -0.21<br>(1.18) | -0.28<br>(1.15) |
| VSN 2 | -0.39<br>(0.65) | -0.37<br>(0.84) | -0.41<br>(1.53) | -0.45<br>(1.65) | -0.37<br>(1.02) | 0.08<br>(0.67) | -0.48<br>(1.11) | -0.26<br>(1.29) |
| VSN 3 | -0.19<br>(1.15) | -0.14<br>(0.91) | 0.10<br>(1.18) | -0.01<br>(1.51) | -0.48<br>(0.80) | -0.31<br>(1.02) | -0.33<br>(0.93) | -0.27<br>(1.03) |
| Stroke | -0.47<br>(0.60) | -0.47<br>(0.59) | -0.48<br>(0.89) | -0.43<br>(1.29) | -0.45<br>(0.96) | -0.14<br>(1.24) | -0.47<br>(1.20) | -0.28<br>(1.43) |
| Healthy | -0.41<br>(0.99) | -0.56<br>(0.89) | -0.48<br>(1.06) | -0.83<br>(0.99) | -0.93<br>(1.18) | -0.56<br>(1.06) | -0.75<br>(0.94) | -0.41<br>(0.89) |

According to the t-tests in the VSN group at pre-rehabilitation (VSN 1), at post-rehabilitation (VSN 2), and at follow-up (VSN 3) no significant deviant minus standard difference potentials emerged. In the Stroke control group, the difference potential was significant for the Left OFF condition at both the left (t(15)=-3.13, p=0.027) and right (t(15)=-3.21, p=0.027) ROIs (Benjamini-Hochberg adjusted values). For the ON events, no significant vMMN emerged in this group. In the Healthy control group, the Left OFF condition was significant at the right ROI (t(16)=-2.57, p=0.040), the Right OFF condition at the left ROI (t(16)=3.26, p=0.013), the Left ON condition at the right ROI (t(16)=-3.46, p=0.013), and the Right ON condition at the left ROI (t(16)=-3.31, p=0.013) (Benjamini-Hochberg adjusted values).

Based on these results, we conducted further analyses with ANOVAs with factors of *Side* (left-, right-side stimulation) x *Stimulus* (deviant, standard) x *ROI* (right, left) in the two control groups. In the Stroke control group for the OFF events, the main effect of *Side* (F(1,15)=4.82, p=0.044, η_p_^2^=0.24), and the main effect of *Stimulus* (F(1,15)=6.23, p=0.024, η_p_^2^=0.29) were significant. In the Healthy control group for the OFF events, the *Side x ROI* interaction (F(1,16)=8.53, p=0.009, η_p_^2^=0.34) and the *Side x Stimulus x ROI* interaction (F(1,16)=6.11, p=0.025, η_p_^2^=0.27) were significant, respectively. According to the Tukey HSD tests, the deviant minus standard difference from left-side stimulation was significant at the right ROI (p = 0.002), while the difference from right-side stimulation was significant at the left ROI (p = 0.006). For the ON events, the main effect of *Stimulus* (F(1,16)=8.69, p=0.009, η_p_^2^=0.35), the *Side x ROI* interaction (F(1,16)=23.88, p=0.0001, η_p_^2^=0.59), and the *Side x Stimulus x ROI* interaction (F(1,16)=21.51, p=0.0002, η_p_^2^=0.57) were significant, respectively. According to the Tukey HSD tests, the deviant minus standard difference from left-side stimulation was significant at both ROIs (p = 0.0001; p = 0.006), and the difference from right-side stimulation was significant at the left ROI (p = 0.0004).

As only the Left OFF condition emerged as significant in both control groups, we then run an ANOVA with factors of *Group* (Stroke, Healthy) x *Stimulus* (deviant, standard) x *ROI* (right, left) to see if there are any differences between these two groups, but there were no significant results with Group as a categorical factor.

## Discussion

In this study we aimed to investigate how early visual processing is involved in the impairments labelled as visuospatial neglect (VSN). This investigation is important because according to influential theories, VSN impairs the attentional system (Corbetta & Shulman, 2011; Bartolomeo et al, 2012), therefore, such accounts expect less deficiency of the brain electric correlates of the automatic (preattentive) processing. We investigated the early exogenous event-related potential components C1 and C2/N1 to task-unrelated stimuli and the signature of automatic detection of violated sequential regulations, the visual mismatch negativity (vMMN). Therefore we compared groups of VSN patients, stroke-without-VSN patients, and healthy age-matched control participants with the application of the OFF-ON method (Sulykos et al., 2017; Csikós et al., 2023).

Concerning the behavioral results, the high performance (hit rate) of the three groups indicated that they were able to attend to the task-related central stimuli, consequently they looked at the middle of the screen, and the ERP-related events stimulated the intended lateral parts of the visual fields. As a sign of general recovery, hit rate in the VSN group showed significant improvement between post-rehabilitation and follow-up, and at which point, their results were no longer different from the two control groups’. This implies the rehabilitation of such nonspatial functions as arousal and alertness, both affected by both stroke and neglect, but their generally increased RT is a sign of the severity of damage.

ON events in the healthy control group elicited the C1 and C2/N1 exogenous ERP components with 60-80 ms, and 120-140 ms latency, respectively. OFF events only elicited the C2/N1 component, and the reason behind the missing C1 for OFF events is probably the sudden ‘lack of’ lines or contours to process, while the ON events mean the reappearance of those parts, thus stimulating the visual field. However, no C1 activity emerged in the VSN group and only approached the significance level in the stroke group. The lack of C1 in the VSN group was obvious in both the initial and later sessions. C1 is generally considered the first cortical visual ERP component, considered as the correlate of pattern onset in V1, described first by Jeffreys (e.g., Jeffreys and Axford 1972). C1 is traditionally considered the initial step of a cascade of processes from V1 up to the ventral and dorsal stream of visual processing. In fact, in earlier studies (e.g. Clark and Hillyard, 1996), no selective attention effects appeared on C1, but according to a more recent meta-analysis (Qin et al., 2022) selective attention modulates C1, i.e., this component is sensitive to top-down effects.

Contrary to the view of V1 (striate) origin of C1, in a recent study we (Csikós et al., 2024) obtained that in the C1 latency range, a fairly large part of the visual cortex was active, including the lateral occipital areas and the lingual cortex. It should be noted that some studies with the aim to localize C1 applied stimuli without contrast onset, such as Gabor patches (Gebodh et al., 2017), or pattern-reversal (Capilla et al., 2016). In a study using checkerboard design, Maier et al. (1987) localized C1 in both the striate and extrastriate cortex, and, applying the onset of alphanumeric characters, Raid et al. (2023) recorded C1 in the extrastriate cortex. In the only study that investigated early visual ERP components in VSN, Di Russo et al., (2008) presented Gabor patches, i.e., stimuli without sharp contours to the four quadrants of the visual field. Contrary to our results, the earliest VSN-related difference appeared after 130 ms, on the N1 components. Besides the highly dissimilar stimulus conditions, the appearance of task-related central stimulation of the present study led to processing differences of the peripheral events.

In the following latency range (C2/N1) we obtained no ERP difference among the three groups, and no difference among the three sessions in the VSN group. In a stage-related model of visual processing with the assumption that the subsequent components represent subsequent stages of processing, i.e., emergence of a later component is contingent upon the processes indicated by a previous one, this is an apparently unexpected result. This result shows a complex relationship between the structures involved in the exogenous components.

While the exogenous ERP components are signatures of the sensitivity of brain structures to various environmental features and endogenous (top-down) effects (like attention), visual mismatch negativity (vMMN) is an index of a specific process, i.e., the automatic detection of violation of sequential regularities of visual input. Being an index of automatic processes, on the basis of theories that connect VSN to attentional processing (Vallar, 1998; Kherkoff, 2001; Corbetta & Shulman, 2011; Bartolomeo et al, 2012), one may expect intact vMMN in this patient group. Contrary to this, we found that no vMMN emerged to either left or right-side stimuli, ON or OFF events in neglect patients.

So far automatic detection of violated sequential regularity in neglect has been investigated only in the auditory modality (MMN). Deouell et al. (2000) and Doricchi et al. (2021) obtained impaired MMN to stimuli contralateral to the damage, but preserved MMN to ipsilateral ones. These results fit the expectation from the lateralized damage, but they also show that even automatic processing was impaired lateral neglect. It should be noted that although Deouell et al. (2000) observed a robust lateral effect for location deviancy, from the 10 participants there were 4 with MMN to left-side deviancy, i.e., to deviants contralateral to the impaired side. As a difference between the results of the two studies, Deouell et al. (2000) reported decreased MMN, whereas in the Doricchi et al. (2021) study no MMN emerged on the contralateral side. It is important to note that both in the Deouell et al. (2000) and the Doricchi et al. (2021) studies, the cognitive demand was low: In the former, participants watched a silent movie, and in the latter study only fixation was requested. Furthermore, in these studies only single oddball sequences were presented. On the contrary, in the present study participants performed a discrimination task, requiring continuous visual attention to the center, and two simultaneous oddball sequences were delivered to the left and right side of the visual field. This difference is not without importance, as here the participants had to have reserves to process two independent sequences, filtrate out the rules governing the stimuli within each sequence, and then process if there was a deviant event violating the established predictions, all the while leaving enough capacity to monitor the other side as well. This is, admittedly, a much more difficult situation for the brain (especially for one after the structural and functional damage caused by the stroke and then neglect), but one that is much more likely to happen in everyday situations.

We suggest that in cases of low cognitive load the limited-capacity attentional system may compensate the impaired automatic system, and accordingly MMN emerged at the contralateral side. However, in case of higher demand (like in the present study) even the non-impaired brain structures are unable to provide such compensation. Moreover, our results come from stimulating a different modality, one that is found impacted more often, or more severely in neglect than hearing, and such difference might easily cause altered results from these aMMN studies. Note, that this assumption preserves the general notion about the attentional damage of unilateral neglect, but the present results indicate the possibility of damage at earlier automatic processes.

With that in mind, one could then suggest that with time elapsed since the initial stroke, and weeks or sometimes months-long rehabilitation, these effects could be mitigated. Unfortunately, the results show that no vMMN emerged in VSN at post-rehabilitation, nor at the 4-months follow-up. Out of these two, the latter results should be treated with more caution as out of the original 17 neglect patients, only 13 returned to participate in that ERP recording session. Although the remaining 4 participants were not able to as a result of severe stroke-related complications, so even for them we can assume the lack of improvement in these processes. Nevertheless, the results from post-rehabilitation still show a lack of improvement in these abilities, suggesting a rather severe and stable deficiency in automatic change detection.

Concerning the lack of vMMN together with the impairment of the C1 components in the VSN group, there are two possible explanations, but both emphasize the damage in automatic processing. First, vMMN emergence depends on the processes indexed by the C1 component. C1 is an ERP component specifically sensitive to pattern onset. (e.g. Jeffreys and Axford, 1972). In fact, in previous studies vMMN emerged without C1 to stimulus offset (Czigler et al., 2019, Sulykos et al., 2017). The other, and more plausible explanation is that the cascade of processes underlying vMMN are compromised in the VSN group. Accepting that vMMN is a signature of automatic processes, the results support the notion that VSN is not only an attention-related syndrome (Vallar, 1998; Kerkhoff, 2001; Corbetta & Shulman, 2011; Bartolomeo et al, 2012).

Nevertheless, this study had some limitations. For one, the number of electrodes. The source of vMMN is dominantly found in the posterior areas of the brain, mostly in the occipital lobe. In that sense, the current design, i.e. 8 electrodes covering that area, and 2 frontal ones should be sufficient for the purpose of this study. While we originally intended to use more, the decision to include only these electrodes resulted from the experience that some of the earliest patients participating became too exhausted during the experiment, thus they could not finish the tasks. In order to avoid that, we chose to shorten the length of the data sampling by limiting the number of electrodes we had to apply. In future research, a denser electrode set, one that equally covers the whole surface of the head would be useful, as that would enable us to also conduct source analyses, similarly to Doricchi et al.’s study (2021), to better understand which parts of the brain are active during the vMMN signal, and to compare the results between healthy and stroke/VSN participants. Another issue could be the number of participants in each group, but the current numbers match those of other studies on average, both in the area of vMMN and neglect research (Di Russo et al., 2008, 2013; File et al., 2017; Czigler et al., 2019).

To summarize the results, this study found that the exogenous component C1 is missing entirely in the VSN group even over time and with rehabilitation, and is insufficiently elicited in the stroke control group, compared to the healthy control group where it emerged as expected. This suggests that even as elementary as the early steps of processing visual stimuli can be deficient in VSN, a lower-level cognitive function than attentional abilities, but one that is prerequisite to the latter. To our other question, that is how automatic change detection, an integrated part of predictive coding changes with and during VSN, the results indicate serious damage compared to healthy participants. VMMN, the signal of error detection, did not emerge in the VSN group. In the case of ON events, where C1 should have been present, the absence of this component indicates that the missing vMMN is not simply the result of deficient change detection, but that even the standard stimuli, the basis of regularity, were not processed properly. Altogether these results point toward the idea that beside attention-related symptoms, the automatic processes of detecting regularities and events violating those regularities also function deficiently in VSN.

